# Tracking Coordinated Cellular Dynamics in Time-Lapse Microscopy with ARCOS.px

**DOI:** 10.1101/2025.03.14.643386

**Authors:** Benjamin Grädel, Lea Brönnimann, Paolo Armando Gagliardi, Lucien Hinderling, Olivier Pertz, Maciej Dobrzyński

**Affiliations:** Institute of Cell Biology, University of Bern, Bern, Switzerland; Graduate School for Cellular and Biomedical Sciences, University of Bern, Bern, Switzerland; Department of Oncology, University of Torino, Torino, Italy

**Keywords:** Cell biology, Image Analysis, Actin Dynamics, Spatial Clustering, Tracking

## Abstract

Subcellular structures like actin waves, podosomes and focal adhesions exhibit dynamic organization that traditional tracking methods struggle to capture. We developed ARCOS.px, an open-source Python package with an interactive napari plugin, to identify and track spatial clusters in microscopy time-lapse images. Unlike conventional approaches that focus on individual particles or bulk intensity changes, ARCOS.px tracks spatial relationships, coordination, and lineage evolution over time. We demonstrate the method by tracking podosome-like structures and RhoA activity waves triggered by growth factors or optogenetics. We revealed how various drugs affect the stability, mobility, and interactions of podosome-like structures in REF52 cells. Additionally, we discovered that actin waves in these cells are followed by active RhoA and Myosin, suggesting feedback mechanisms that recruit active RhoA at their trailing edge. ARCOS.px can be applied to diverse research contexts requiring the tracking of discontinuous yet dynamically associated regions.

## Introduction

Spatiotemporal organisation of signalling networks enables cells to coordinate complex behaviours such as cell polarization, motility, as well as cell-cell interactions during collective cell migration or transmission of mechanical forces between cells. The self-organizing properties of signalling networks allow cells to rapidly reorganize their internal architecture and effect robust responses to external stimuli. A striking example of such self-organization are patterns of small GTPases and cytoskeletal components, which emerge through reaction-diffusion of signalling molecules coupled with mechanical feedback interactions. The Rho family of small GTPases, in particular, can spontaneously form diverse spatiotemporal patterns in the cell cortex, ranging from static clusters and oscillatory pulses to propagating waves (1).

Similar self-organizing principles govern the assembly of dense actin structures, where nucleation factors, crosslinkers, and regulatory proteins coordinate to create dynamic architectures essential for cell movement. These include lamellipodial networks at the leading edge, contractile stress fibres, and focal adhesion complexes that anchor the cell to its substrate. The spatiotemporal coordination between these structures enables the cyclic processes of protrusion, adhesion, and retraction that drive directional cell migration (2).

Beyond Rho GTPases and actin dynamics, numerous signalling systems exhibit spatiotemporal behaviours at the single-cell level. Calcium signalling displays intricate spatial patterns and travelling waves that can modulate cytoskeletal dynamics and cellular mechanics (3). Phosphoinositide lipids form dynamic membrane domains that help establish and regulate protein recruitment to initiate downstream signalling cascades (4). Even metabolic enzymes can organize into dynamic condensates that influence local reaction rates and signal propagation (5).

Automatic identification and tracking of such coordinated dynamic events from time-lapse imaging data is still challenging, yet essential for further understanding the coupling between signalling and mechanics. Here, we use the term *coordinated dynamic events* to describe phenomena such as actin or RhoA waves that arise from individual components in proximity and influence one another’s formation and progression over time. While some methods provide information on a pixel-level scale for understanding structural characteristics of a system, they lack direct temporal information. For example, OrientationJ (6) extracts structural features from images but doesn’t incorporate tracking capabilities. On the other hand, tracking in cell biology is mainly suited for segmenting cells or their nuclei, which are then tracked across time frames based on objects’ centroids using nearest neighbour or linear-assignment strategies (7, 8), or recent machine learning approaches (9).

In this work, we present ARCOS.px, a computational method to track coordinated dynamic cellular events directly in timelapse microscopy images without needing prior instance segmentation. ARCOS.px builds on our earlier method, AR-COS (Automatic Recognition of Collective Signalling) (10), which was only suitable for tracking collective events from already segmented objects, such as cell nuclei, with an associated measurement. Here, we rewrite and optimise the algorithm to handle raster images, which enabled us to identify and track spatial clusters of pixels that represent activity of subcellular signalling or emergence of dynamic cell structures. Pixel-level tracking allows handling of discontinuous and irregularly shaped regions. We introduce several new features: (i) tracking lineages in phenomena involving merges and splits of spatial clusters, (ii) motion prediction to aid tracking of moving clusters, (iii) alternative spatial clustering, e.g., HDBSCAN, and frame-to-frame linking methods, e.g., optimal transport, (iv) improved memory management to handle large raster images, and (v) *lazy* processing, which makes our framework suitable for online image processing in smart microscopy workflows. Our tool is available as a Python package (11), and we provide a GUI in a dedicated napari plugin (12). It applies to various biological contexts, and fills a gap in current bioimage analysis methodologies by enabling pixel-level event tracking.

## Results

### Algorithm Overview

The algorithm takes a binary image as input, spatially clusters active pixels, and tracks these clusters over time (Figs. 1A and S1A). Image binarisation can be performed either using simple thresholding, machine learningbased pixel classification (13, 14) or any other method that generates a binarised output. The choice of binarisation method depends on the data, ensuring that regions of interest that exhibit spatially correlated dynamic activity are accurately identified and separated from the background. The clustering step utilises the DBSCAN algorithm (15, 16) parametrised by the neighbourhood radius *ε* and a minimal size of the cluster. Alternatively, the user can opt for HDBSCAN (16, 17), which is more flexible and requires less parameter tuning than the default DBSCAN. HDBSCAN is recommended when finding a single *ε* is difficult, e.g., when clusters of varying densities and sizes are tracked. However, HDBSCAN may perform poorly on noisy images, as it will attempt to identify clusters in such data.

**Fig. 1:**
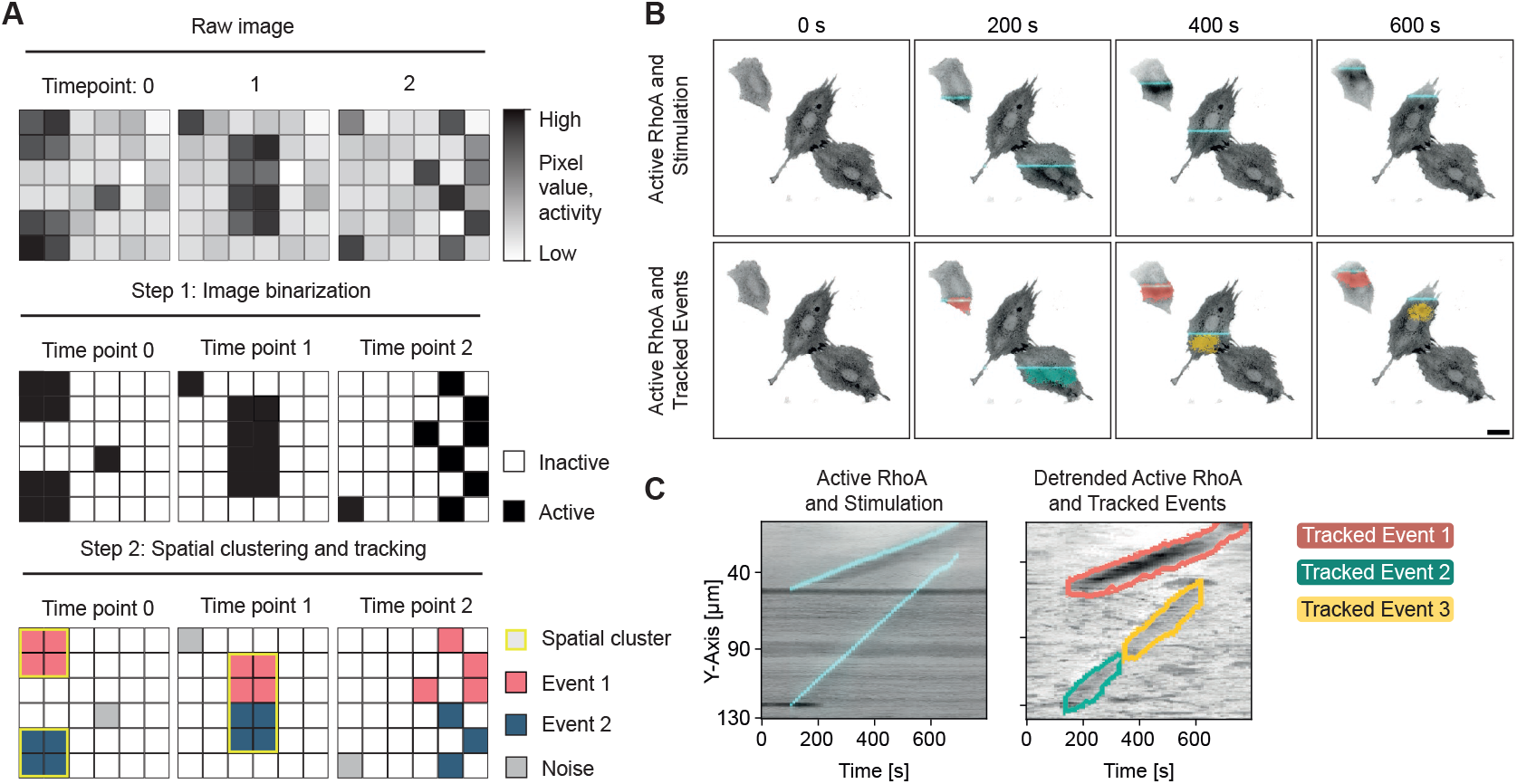
The algorithm. (A) Algorithm scheme. Top: a greyscale image with two regions travelling through the frames. Middle: a binarized activity image used as input for ARCOS.px. Bottom: ARCOS.px output with a label image of tracked dynamic events. Yellow outlines indicate individual spatial clusters identified by DBSCAN. (B) Optogenetic induction of RhoA activity wave in REF52 cells expressing a RhoA activity sensor 2×rGBD and the optogenetic actuator optoLARG as a biological ground truth model to test the algorithm. Top: cells are stimulated by a horizontal line of blue light travelling along the vertical image axis. Bottom: region of active RhoA following the line stimulation, as tracked using ARCOS.px. Scale bar: 20*µm*. (C) Maximum intensity projection along the X-axis of the image stack. Left: a projection of the raw image and stimulation, indicated by a blue line. Right: a projection of background-removed image with detected events, as in panel B.

Clustering is followed by tracking individual pixels within clusters of the current frame to clusters already identified in a previous (or several previous) frame(s). The depth of temporal linking of clusters between frames is set by the *n*_prev_ parameter. Linking is performed by the nearest neighbour assignment, meaning the algorithm iteratively assigns pixels to an event based on their proximity set by the *ε*_prev_ radius. It may happen that pixels in a cluster from a current time frame are linked based on their proximity to two different clusters in the previous time point. Then, the pixels inherit respective cluster IDs from those previous clusters. This case is illustrated in Fig. 1A, where pixels in a single cluster at time point 1 are associated, based on *ε*_prev_, to two already existing clusters at time point 0: event 1 (red) and event 2 (blue).

The ARCOS.px Python library also provides a linking strategy that considers the problem as unbalanced optimal transport (18, 19), using the *sinkhorn* algorithm (20). The advantage of this linking strategy is that it provides an optimal global solution to linking track segments, as opposed to brute force assignment of pixels based only on nearest neighbours. We use the unbalanced version of this algorithm because the number of pixels in a cluster changes between time frames. Furthermore, a linear motion predictor can estimate a future position of a cluster in the next frame based on *n* previous frames, thereby enhancing the overall tracking performance.

### Pre-processing

Image pre-processing can further improve tracking and feature extraction performance. For example, we recommend temporal filtering to remove static image features such as background objects that are not part of dynamic events. Methods range from simple rolling subtraction, rolling filters, or dedicated background subtraction algorithms as implemented in the OpenCV library (21). This is especially relevant for tracking of dynamic features in images with a low signal-to-noise ratio.

Preprocessing can also include downscaling or image binning, i.e., merging square image patches to the minimally required resolution, where the features are still clearly visible, but without redundant information that would slow down the tracking algorithm. Such a step improves tracking accuracy and the overall computation speed.

### Availability

Raw imaging data with annotations are available on the Bioimage Archive (22). ARCOS.px code is available as a Python package through PyPI and conda-forge, with an API to apply the method directly to raster images (11). Documentation and notebooks required to recreate this analysis is available on GitHub (23). ARCOS.px can also be used as a dedicated plugin (Fig. S1B) (12) in an interactive interface of the napari image viewer (24).

### Testing the algorithm

To test the limits of ARCOS.px, we applied it to simulated ground truth data, and then evaluated it in a biological system with known wave behaviour.

To generate ground truth tracking data, we implemented acellular automaton as defined in Fig. S2 and Materials and

Methods. With this framework, we simulated circular and directional/travelling waves, as well as target and chaotic patterns (Fig. S3A, Video S1). We also added Gaussian noise to the images to see how the algorithm would perform in suboptimal acquisition conditions (Vids. S2–S5).

By comparing waves detected by ARCOS.px to the ground truth from simulated patterns, we quantified tracking accuracy using two metrics: Multiple Object Tracking Precision (MOTP) and Accuracy (MOTA) (see Eqs. 4,5 in Materials and Methods) (25). All types of patterns were correctly segmented and tracked; however, noise affected the tracking differently depending on the pattern. At a moderate noise level of 0 dB, which corresponds to equal powers of noise and the signal, tracking of circular patterns (Fig. S3B) was still acceptable as evidenced by the relatively low number of false positives and missed ground truth detections, and the lack of switches in track IDs (Fig. S3C). The tracking accuracy metric MOTA = 0.5 compared to 0.97 in the absence of noise (1 being the maximum), and tracking precision MOTP = 0.07 compared to 0.02 without added noise (0 being the minimum). Despite higher spatial complexity, tracking of simulated directional and target waves was only slightly more susceptible to noise compared to circular patterns (Fig. S3D).

To supplement simulated tests, we evaluated ARCOS.px on a biological system with synthetically induced waves. We used rat embryonic fibroblasts, REF52, cells with stably transfected optogenetic actuator, optoLARG, to reversibly manipulate RhoA activity in single living cells (26, 27), and the RhoA biosensor, 2×rGBD (28). We used a digital micromirror device, which can illuminate a field of view with spatial patterns defined on an 800×600-element grid, to synthetically generate a wave of RhoA activity. We projected a line of blue light that travels across the cells’ surface, with a velocity determined by the vertical axis length of each cell. This light stimulation transiently activates optoLARG, which is recruited locally to the plasma membrane, thus activating RhoA with a short time delay. As expected, in all three cells acquired in the field of view, we induced a region of RhoA activity trailing behind the moving line (Fig. 1B and Video S6).

As the pre-processing step, we detrended images using a rolling median filter, and downscaled the spatial dimensions by a factor of 4 to speed up the calculations. A fixed threshold was used to identify activity regions. ARCOS.px successfully tracked RhoA activity clusters, despite them being discontinuous, dynamically changing their shape, and having varying fluorescence intensity levels between time frames (Fig. 1C). This experiment reassured us in ARCOS.px’s ability to detect and track subtle intra-cellular events over time, even under low signal-to-noise ratio in our images.

### Tracking Dense Actin Structures

Treatment of REF52 cells with platelet-derived growth factor (PDGF) leads to assembly of dense actin structures, similar to podosomes. In a previous study (29), podosome-like structures (PLSs) in REF52 cells were found to contribute to maintaining cell polarization during migration. Podosomes are dynamic actin structures, mostly composed of branched F-actin, surrounded by a ring of adhesion molecules. They fuse, fission, and display pulsating oscillations (30, 31). We find that classic methods that identify a centre of mass or a bounding box struggle with segmenting and tracking such events. In contrast, ARCOS.px offers a notable advantage by tracking discontinuous, dynamically changing spatial clusters of pixels, which enables capturing the dynamics of fusion events and preserving contributions of the original structures. The pixel-based tracking approach used in ARCOS.px allows for detailed quantification of podosome lifetime, movement, intensity and correlation with other markers. Our implementation of lineage tracking (see Materials and Methods) allows for precise quantification of merges and splits, providing new quantitative insights into PLS formation dynamics. Indeed, with ARCOS.px we were able to track PLSs (Fig. 2A–C and Video S7), their aggregation and fission (Fig. 2D).

**Fig. 2:**
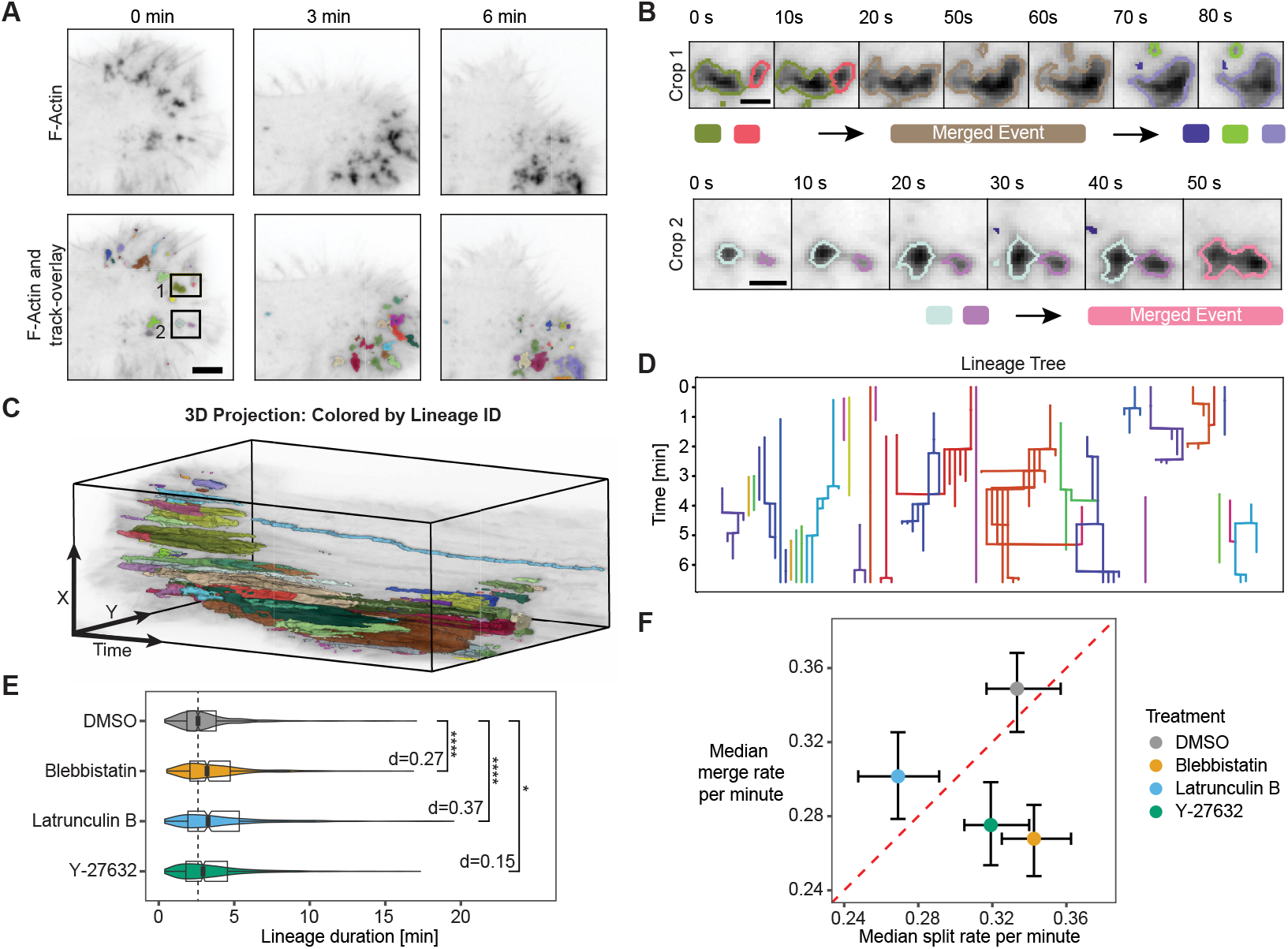
Tracking dense actin structures in REF52 cells. (A) REF52 cells expressing the F-actin marker Lifeact imaged using 100x magnification using a widefield microscope. Top: excerpts from the time-lapse; bottom: overlaid output generated by ARCOS.px. Distinct colours mark individual events. Scale bar: 5*µm* (B) Crops of two selected events from A. Scale bar: 2*µm*. (C) 3D representation of the raw input with tracked events as overlay, coloured by lineage ID. (D) Representation of the PLS lineage tree corresponding to A showing only lineages longer than 30 seconds. Merges occur where branches combine; splits occur where branches diverge from a single root. Colours indicate lineages. (E) Changes in lineage duration in response to drug treatments. Selected lineages, *N ∈* [524, 968] per condition, last longer than 16 *s* (4 frames), with at least one merge and/or split event, do not merge with a bigger lineage. We used a two-sample Wilcoxon test; ****: *p ≤* 0.0001, *: *p ≤* 0.05. d: Cohen’s d effect size. (F) Changes in PLS merge vs. split rate in response to drug treatments. Selected lineages, *N ∈* [1067, 1615] per condition, last longer than 16 *s*, with at least one merge and/or split event. Error bars indicate 95% CI for the median calculated from bootstrapping, *N* = 10*k*. The red dashed line indicates an equal split and merge rate.

To better understand podosome dynamics and regulation, we perturbed PDGF-treated REF52 cells with the myosin inhibitor blebbistatin (20*µM*), an inhibitor of actin polymerisation latrunculin B (0.3*µM*), and ROCK inhibitor Y-27632 (40*µM*). When treated with latrunculin B, the percentage of PLSs without merges or splits increased to 90% from 83% in the DMSO control (*p ≤* 0.0001; Fig. S4A). The PLS lineages that did merge or split under this treatment lasted longer (3.3 vs. 2.6 min in DMSO control; *p ≤* 0.0001; Cohen’s *d* = 0.4), suggesting that blocking actin polymerization creates a more static PLS state (Fig. 2E). With reduced polymerization, podosomes become more stable but less dynamic, which is consistent with earlier observations that showed increased podosome lifetime after inhibiting actin polymerization (32). This reduced remodelling is also evidenced by lower rates of PLS merges and splits under latrunculin B treatment (Fig. 2F).

Cells treated with blebbistatin and Y-27632 showed a significant decrease in median PLS merge rates, *≈* 0.27 min^*−*1^, CI 95% [0.25, 0.29], compared to 0.35 min^*−*1^, CI 95% [0.33, 0.37] for DMSO, while median split rates, *≈* 0.33 min^*−*1^, CI 95% [0.30, 0.36], remained similar to control cells. These rates were calculated only for lineages with at least one merge or split event, as short-lived PLSs do not last long enough for fusion or fission (Fig. S4B). Interestingly, despite blebbistatin and Y-27632 decreasing merge rates, both inhibitors slightly increased PLS lineage lifetime from 2.6 min for DMSO to 3.2 min (*p ≤* 0.0001) for blebbistatin and 2.9 min (*p ≤* 0.05) for Y-27632 (Fig. 2E). This may be explained by multiple progeny structures forming after PLS splitting, which then create many fragmented but persistent PLSs that do not merge, leading to slightly longer lineage duration despite splitting. Video S8 shows this increased PLS fragmentation in blebbistatin and Y-27632-treated cells compared to latrunculin B treatment.

Together, these findings demonstrate that actin polymerization dynamics and myosin II function play distinct yet complementary roles in regulating podosome fusion, fission, and stability. Actin turnover, here reduced by latrunculin B treatment, appears to primarily drive the dynamic aspects of remodelling, affecting how frequently podosomes merge or split. By contrast, myosin II helps maintain the structural cohesion of podosome networks, so inhibiting its activity (via blebbistatin or Y-27632) weakens the forces and crosslinking that hold podosomes together, leading to more frequent fragmentation and fewer fusion events.

### Actin polymerization waves are trailed by RhoA activity and Myosin

Actin can self-organize into propagating waves in neurons, keratinocytes, fibroblasts and other cellular systems (33–36). These waves are commonly generated through activator-inhibitor systems. An activator promotes actin assembly at the leading edge, while an inactivator, typically operating with a temporal delay, facilitates actin disassembly at the trailing edge. This coordinated process orchestrates wave-like propagation across the cell cortex, as described in recent reviews (1, 37, 38). In REF52 cells, such actin polymerisation waves are observed in combination with PLSs triggered by treating REF52 fibroblasts with PDGF. To understand what causes the propagation and collapse of these polymerization waves in this system, we performed imaging of F-actin using the Lifeact reporter (39), RhoA activity using the 2xrGBD reporter (28), and myosin II light-chain (MLC) using total internal reflection (TIRF) microscopy (Fig. 3A and Video S9). We observed that the different fluorescence signals colocalised in these waves. However, from kymographs alone, the temporal order of signals within individual waves is not apparent (Fig. 3B). Using our pipeline, we could accurately quantify delays between the individual signals.

**Fig. 3:**
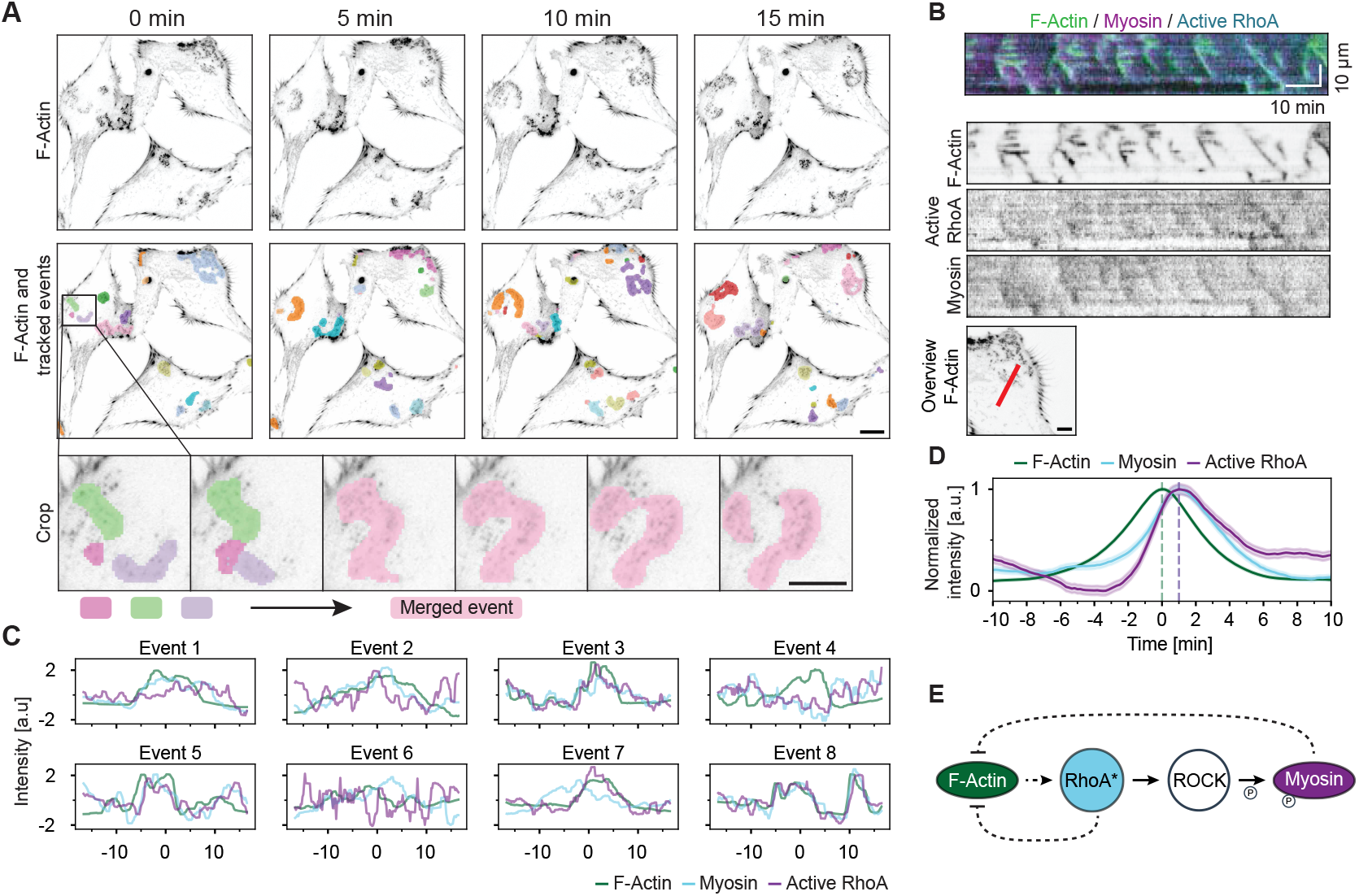
Tracking wave-like dynamics in REF52 cells. (A) Lifeact in REF52 cells imaged with TIRF microscopy at a 10*s* interval. Tracked events are marked with distinct colours. The inset shows one event at higher temporal resolution. Scale bars: 25*µm* in the main panel, 5*µm* in the inset. (B) Kymograph of wave events for F-actin, Myosin II Light Chain (MLC) and active RhoA (2xrGBD) along the red line indicated in the overview panel. The scale bar in overview: 10*µm*. (C) Intensity measurements for a sample of 8 out of *≈* 5000 windows from 175 events lasting longer than 200*s*. (D) Averaged intensities of *≈* 5000 windows such as those in C. The dashed line indicates the peak value: Active RhoA (2xrGBD) and Myosin peaked at *s* F-actin. Shaded regions indicate 95% Confidence Interval. (E) Proposed circuit with hypothetical feedback mechanisms that could generate the observed wave propagation, indicated with dashed lines.

To prepare images for the analysis, we removed static image features using temporal filtering. To analyse temporal shifts between RhoA activity, F-actin and myosin, we need tracked masks for every wave event, which we obtain using ARCOS.px. We then extracted peak-aligned kymographs from the three imaged channels for individual waves (see Materials and Methods). After averaging over individual intensity profiles from Fig. 3C, we obtained the intensity curves shown in Fig. 3D. From this, we inferred that waves are led by actin and trailed by active RhoA and MLC with a delay of *≈* 60*s*. This suggests that these actin polymerisation waves recruit an activator of RhoA. In line with the known function of RhoA and its effector ROCK (40), we see an increase in RhoA activity accompanied by increased MLC. Assuming an activator-inhibitor system, one possibility would be an activation via either a nucleator of F-actin, F-actin itself, or actin-associated protein to RhoA. Subsequently, RhoA or one of its downstream targets such as phosphorylated myosin could act as a negative regulator of F-Actin (Fig. 3E), potentially in a similar manner as described in Rao *et al*. (41).

Using ARCOS.px, we extracted new high-quality information regarding components within actin polymerisation waves, which let us generate new hypotheses about the underlying mechanism of wave emergence and propagation.

### Tracking integrity of epithelial monolayer

To demonstrate ARCOS.px on a tissue scale phenomenon, we quantified loss of epithelial integrity by measuring the area of holes formed in a monolayer of MCF10A WT cells due to dying cells in response to treatment with a chemotherapeutic drug, doxorubicin (Fig. 4A and Video S10). Growing holes have an undefined, rapidly changing shape, with cases of hole fusion. Cells were stably transfected with biosensors for ERK and AKT activity, and a nuclear marker (see Materials and Methods). We segmented holes based on averaged fluorescence intensity from ERK and AKT channels with Ilastik’s pixel classifier (13), which provided binarised input for cluster tracking with ARCOS.px. Consistent with our earlier observations, high doses of doxorubicin increased apoptosis and rapid hole formation that reached almost 50% of the field of view at 24h after treatment with 5*µM* of doxorubicin (42) (Fig. 4B). With our pipeline, we extracted information about the dynamics of individual hole formation (Fig. 4C).

**Fig. 4:**
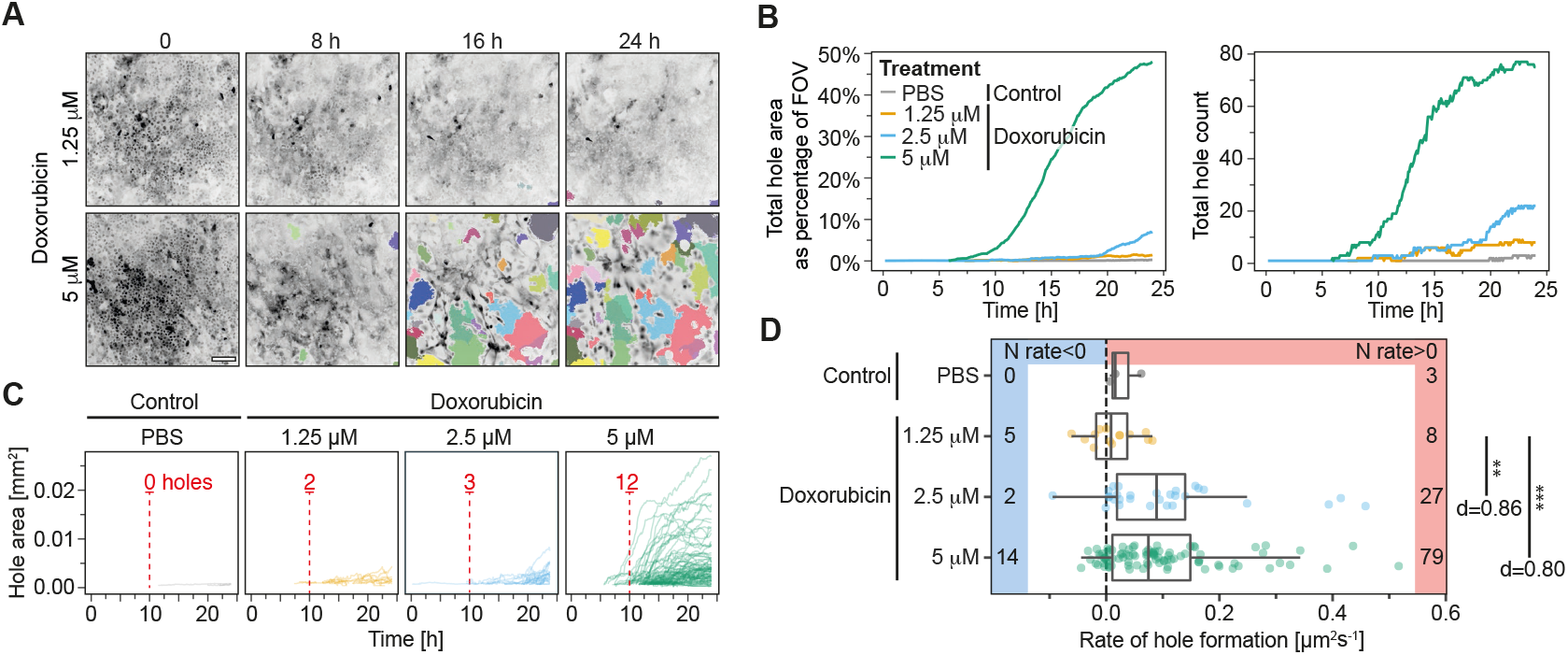
Tracking hole formation in the MCF10A epithelial monolayer. (A) Hole formation in response to low and high dosage of doxorubicin applied at time 0. Shown is ERK activity quantified by the ERK-KTR translocation reporter, as described earlier (42). Holes, indicated by coloured regions, were initially segmented using Ilastik (13) and then tracked using ARCOS.px. Scale bar: 100*µm*. (B) Dynamics of hole formation in response to doxorubicin collated from two fields of view and quantified as a percentage of the FOV area and as the total hole count. (C) Growth of individual holes in the monolayer. (D) Rate of individual hole formation, calculated as the ratio of each hole’s total area increment and its lifetime. We used a two-sample Wilcoxon test; ***: *p ≤* 0.001, **: *p ≤* 0.01. d: Cohen’s d effect size. The number of holes with the negative (shrinking) and positive (growing) formation rate are provided on the margins.

We find that the median rate of individual hole growth is 0.09*µm*^2^*/s*, CI 95% [0.02, 0.12], for 2.5*µM* and 0.07*µm*^2^*/s*, CI 95% [0.05, 0.09], for 5*µM* doxorubicin, with the majority of holes having a positive rate, and thus growing (Fig. 4D). However, the onset of the rapid hole formation, and hence the collapse of the epithelial monolayer, is *≈* 10*h* earlier for the highest dosage as evidenced by the rapid onset of the total hole area growth and the hole number at 10h posttreatment (Fig. 4B,C). By tracking individual holes, we find that the lowest treatment (1.25*µM*) induced 5 slowly shrinking and 8 slowly growing holes during 24h of acquisition, resulting in minimal epithelial damage with only 1.5% of the total area covered by holes at 24h post-treatment. Conversely, the majority of holes for 5*µM* doxorubicin displayed high, positive growth rates, leading to a 48% hole coverage at 24h (Fig. 4D).

## Limitations

ARCOS.px relies on image binarisation to help identify regions of activity. Binarised raster images are then spatially clustered using DBSCAN, which introduces dependencies on the clustering distance, *ε*, and minimum cluster size. We provide a function that estimates *ε* based on the nearest neighbour distance, however, these parameters often need to be determined on a case-by-case basis and the guidelines only provide a rough starting point. An alternative, HDBSCAN, doesn’t require setting a maximum distance but could detect false positives in case of noisy data. Additionally, high object density can cause tracking inaccuracies, artefacts, and decrease the processing speed. In Fig. S3, we evaluated our algorithm on simulated noisy time-lapses and in our previous work (10) we further quantified the performance for a range of object densities.

## Conclusions

We present a computational method, ARCOS.px, to track emergent dynamic cellular events. Classic segmentation frameworks involve labelling directly connected regions, which are then identified as objects and represented by object features such as centroids or bounding boxes. Our densitybased clustering creates clear segmentation masks of noncontiguous objects, which enabled us to track (i) prototypical RhoA activity waves (Fig. 1B, C), (ii) the effect of drug treatments on dynamics of podosome-like structures (Fig. 2), (iii) actin polymerization waves (Fig. 3), and (iv) hole formation in a drug-treated epithelial monolayer (Fig. 4).

In Fig. 2, we gained novel biological insights into podosome regulation and dynamics. Latrunculin B treatment of REF52 cells decreases podosome splitting and merging rates and prolongs podosome lineage lifetime in the subpopulation of podosomes that undergoes splits and merges. This underscores the critical role of actin polymerization in podosome dynamics and turnover. Prior studies demonstrated that cytochalasin D, also an inhibitor of F-actin polymerisation like latrunculin B, treatment decreased podosome-mediated protrusive force and oscillation amplitude (43), and increased lifetime (32) implying that inhibited actin turnover interferes with the normal cycle of core expansion and collapse. We now find that inhibition of F-actin polymerization also leads to increased podosome lifetimes, possibly by altering the splitting and merging rate of those structures.

Our quantification indicates that blebbistatin and Y-27632 treatments reduce the merge rate compared to DMSO control. This suggests that myosin II–function and ROCK-mediated signaling influence the fusion-fission balance. ARCOS.px enabled us to measure podosome splitand merge dynamics as well as lifetime statistics.

In Fig. 3, we measured delays between different fluorescent reporters, enabling us to propose a hypothetical model of Factin wave propagation in REF52 cell stimulated with PDGF. Although further experiments and simulations are required to understand the exact mechanism at play, ARCOS.px provided us with a method to easily extract dynamic information from noisy images.

A study by Martin *et al*. (29) demonstrated that PLSs orchestrated fibroblast migration and contributed significantly to maintaining cell polarity in these cells. They showed that these PLSs were associated with a zone of low RhoA activity, and that phosphorylated MLC accumulated behind the podosome-zone. Our analysis confirmed these results as we see podosome waves trailed by active RhoA and myosin II, not only at the cell periphery but also within the cell. In cases where podosome waves hit the edge of the cells, an outwards directed membrane deformation can be observed.

In the model system described in (29), one could speculate that a stable wave assembles at the front, which continuously pushes the cell forward.

In Fig. 4, we quantified how epithelial barrier integrity deteriorates across different doxorubicin concentrations. The pattern and timing of hole formation may reveal underlying biological mechanisms, e.g., whether holes form randomly or at specific structural weak points. With sufficient data across conditions, one could develop models that predict tissue damage based on early formation patterns. The statistics gathered with our method could be used to compare candidate compounds or combination therapies that might mitigate doxorubicin-induced tissue damage and could help predict individual sensitivity to treatments.

ARCOS.px enables researchers to quantify previously difficult-to-measure dynamic properties of subcellular structures, which makes it unique for studying dynamic, self-organizing cellular systems. Due to its modular architecture, ARCOS.px can easily utilize custom linking and clustering functions. For example, GPU-accelerated density-based clustering from the cuML library could improve the speed of large datasets. ARCOS.px integrates into the napari plugin ecosystem and can be combined with already available napari plugins. This enables the construction of complex image analysis workflows. Already in Fig. 2, to segment podosomes we used ARCOS.px in combination with another napari plugin, a pixel classifier convpaint (14). Also, filters and initial segmentation steps can be performed with the napari-skimage or napari-assistant plugins. We are committed to supporting the library further with fixes and feature upgrades. Our extensive documentation and code can be used as a tutorial to apply ARCOS.px to other systems.

## Materials and Methods

### Simulated waves

To benchmark ARCOS.px, we implemented a two-dimensional cellular automaton to simulate wave propagation across an excitable medium. The simulation tracks cells on four primary grids: an activity grid recording cell states, a refractory grid monitoring recovery periods, a lifetime grid tracking continuous activation duration, and a wave ID grid for identifying distinct wavefronts. Updates of cell states follow probabilistic rules. Active cells may deactivate with a probability that increases with their lifetime, according to:

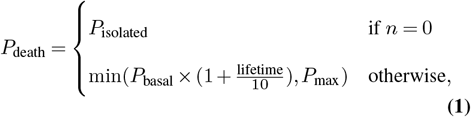

where *n* is the number of active neighbours, *P*_isolated_ is the death probability for isolated cells, *P*_basal_ is the basal death probability, *P*_max_ is the maximum death probability, and lifetime is the duration of cell’s activity.

Inactive, non-refractory cells can activate through neighbour influence with probability:

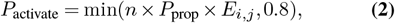

where *n* is the number of active neighbours, *P*_prop_ is the propagation probability, and *E*_*i*,*j*_ is the local excitability that influences each cell’s activation probability individually. Cells may also activate spontaneously with probability *P*_spontaneous_ = *P*_wave_ *× E*_*i*,*j*_, where *P*_wave_ is the wave formation probability.

Each cell undergoes a refractory period after deactivation, during which it cannot be reactivated. Waves are tracked by propagating their identifiers, which allows analysis of individual wavefront dynamics across the grid over time. When a cell becomes active due to its neighbours, it inherits the most common wave ID among those neighbours. If a cell activates without neighbours or all neighbour wave IDs are inactive, it is assigned a new unique wave ID.

We added Gaussian noise with intensity dependent on the signal-to-noise ratio (SNR) to evaluate ARCOS.px under different signal-to-noise conditions. For an input image *I*, the noisy image *I*_*n*_ is given by:

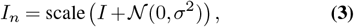

where *σ*^2^ = *P*_signal_*/*(10^SNR*/*10^), *P*_signal_ = 𝔼 [*I*^2^], and scale() ensures the output is in the range [0, 2^16^ *−* 1].

For each wave pattern, we set different parameters to generate specific patterns. The code to run the simulations and reproduce benchmark results is available on GitHub (23).

### Tracker performance metrics

Evaluation of tracking performance was performed with the py-motmetrics python library. We used two metrics: Multiple Object Tracking Accuracy (MOTA) and Multiple Object Tracking Precision (MOTP) (25). MOTA measures the overall accuracy of the tracker and detection:

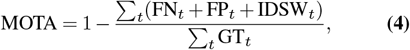

where *t* is the frame index, FN is the number of false negatives, FP is the number of false positives, and GT is the number of ground truth objects. IDSW is a count of identity switches – mismatch errors counted if a ground truth target is matched to a new track. In our benchmarks, we report MOTA on the (*−∞*, 1] range, with 1 being the ideal scenario without false positives, negatives, and track ID switches. Negative MOTA values occur when the number of errors made by the tracker exceeds the number of objects in the FOV.

MOTP describes the average overlap between all correctly matched hypotheses and their respective objects:

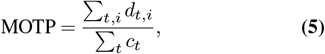

where *d*_*t*,*i*_ = 1 *−* IoU_*t*,*i*_ is the distance between the localization of objects in the ground truth and the detection output *i*, where IoU is the intersection over union, i.e., an overlap area calculated between bounding boxes of the prediction *i* and the ground truth. *c*_*t*_ denotes the number of matches in frame *t*. A match is counted if the overlap exceeds the threshold, which in our case was *t*_*d*_ = 0.5. For a perfect match and a full overlap between objects MOTP = 0.

### Cell culture and Microscopy

#### REF52 cells

Cells were grown at 37°C and 5% CO2 in Dulbecco’s Modified Eagle Medium (DMEM) with 4.5 g/L glucose, 10% 4mM L-Glutamine, and 100 U/mL penicillin/streptomycin. Imaging was performed in FLuorobrite DMEM, supplemented with 0.5% FBS, 0.5% BSA, 10% 4mM L-Glutamine, and 100 U/mL penicillin/streptomycin with cells seeded in a glass-bottom well plate coated with fibronectin.

#### Optogenetic experiments

REF52 cells were stably transfected using optoLARG and 2xrGBD-dTomato and imaged on a Nikon Eclipse Ti Microscope using a 60x Plan Apo VC objective (NA 1.4). Images were acquired on Andor Zyla 4.2 with 2x2 binning, at a 10s interval.

#### PLS tracking and lineage analysis

REF52 cells were stably transfected with Lifeact-mCherry (Fig. 2A–D), imaged with a 100x Plan Apo TIRF objective (NA 1.49), or Lifeact-mNeonGreen (Fig. 2E,F), respectively, imaged with a 60x Plan Apo TIRF objective (NA 1.49) and 1.5 intermediate magnification. Cells were seeded in serum-starved medium containing 50 ng/mL PDGF in a glass-bottom well plate coated with fibronectin.

#### PLS drug perturbations

Forty-eight hours after PDGF addition, REF52 cells were preincubated with either blebbistatin (*≈* 3.5*h*, diluted 1:250 to a final concentration of 20*µM*), Y-27632 (*≈* 3.5*h*, diluted 1:250 to a final concentration of 40 *µ*M), latrunculin B (*≈* 30*min*, diluted 1:80000 to a final concentration of 0.3 *µ*M), or DMSO (*≈* 3.5*h*, diluted 1:250). For each condition, 10 fields of view were sequentially acquired. Images were acquired on a Nikon Eclipse Ti Microscope equipped with a prime 95B sCMOS camera, with binning set to 1×1.

#### TIRF microscopy

REF52 cells were stably transfected with Lifeact-mNeonGreen, 2xrGBD-dTomato, and Myosin II light chain-miRFP703 and imaged in Cellvis glass bottom plates using a Nikon Eclipse Ti Microscope equipped with an Ilas2 TIRF system using an Apo TIRF 60x oil immersion objective (NA 1.49). The system was equipped with a prime 95B sCMOS camera set to 2x2 binning, and images were acquired at 10s intervals. 50 ng/mL PDGF was added 24 to 48 hours before the experiment started.

#### Hole tracking in epithelial monolayer

MCF10A WT cells were grown at 37°C and 5% CO2 in DMEM:F12 supplemented with 5% horse serum, 20 ng/mL recombinant human EGF 10 *µ*g/mL insulin, 0.5 mg/mL hydrocortisone and P/S. Experiments were carried out in starvation medium composed of DMEM:F12 supplemented with 0.3% BSA (Sigma-Aldrich/Merck), 0.5 mg/mL hydrocortisone (SigmaAldrich/Merck), 200 U/mL penicillin and 200 *µ*g/mL streptomycin. For hole tracking, cells were stably transfected with ERK-KTR-mTurquoise2 (for ERK activity), FoxO3amNeongreen (for AKT activity) and H2B-miRFP703 (for nuclear marker) with PiggyBac plasmids as described in Gagliardi *et al*. (42). Cells were cultured overnight in starving medium and imaged for 24h at 5-minute intervals with a Plan Apo air 20x (NA 0.8) objective.

#### Image analysis

Image analysis was performed using the Python scientific ecosystem. The key libraries used are numpy (44), scipy (45), scikit-image (46), scikit-learn (16) matplotlib (47) and napari (24).

In most cases, we preprocessed microscopy images by subtracting a rolling median filter applied over each pixel across time, which served as the background subtraction method. Images were then binned using an n-by-n median filter to reduce the number of pixels tracked, where the full resolution was not required. We used *n* = 2 or 4.

#### Split and Merge Detection

Potential splits and merges are identified and tracked over time to ensure they represent stable topological changes rather than transient fluctuations. A potential merge is identified when multiple propagated cluster IDs in the current frame map to a single cluster identified by the clustering algorithm. Conversely, a potential split occurs when a single linked cluster ID maps to multiple original clusters. A split or merge is stable if the new state lasts for a predefined number of frames. To detect this, each cluster gets assigned a stability measure over time, *S*_cl_(*t*), which quantifies the number of frames *f* from a set ℱ where the new configuration was observed in a time window of *w* frames:

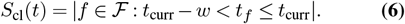

A split or merge is stable when the duration of the new configuration exceeds the cluster’s stability threshold, *w, S*_cl_(*t*) *≥ w*. Furthermore, splits are valid when the size of each new sub-cluster *C*_sub_ exceeds a minimum threshold, |*C*_sub_| *≥ k × n*_min_, where *k* is a multiplier (typically *≥* 1), and *n*_min_ is the minimum cluster size. Similarly, merges are valid when the resulting merged cluster exceeds the minimum size, |*C*_merged_| *≥ n*_min_. Stable merges and splits are recorded as edges and nodes, accessible in the output. When two or more lineages merge, the progeny lineage inherits its lineage id from the parental lineage with the largest area. When a lineage splits, all progeny inherit their lineage id from the parent.

#### Lineage Analysis

Raw images were cropped to 300×300 pixels around the region of interest. Convpaint (14) was then used to segment foreground from background, and the resulting binary images were passed to ARCOS.px. The stability threshold *w* = 5 from Eq. 6 was used for split and merge detection. All processing scripts are available on GitHub (23).

#### Windowed, Peak-Aligned Intensity Extraction

We developed an automated pipeline that extracts local, peakaligned kymographs of fluorescence intensity around events, thus allowing us to pool multiple events and capture general trends in signal dynamics. The data and code used for reproducing all figures are available on GitHub (23).

For each event, at its every time point *m*, we fix the event’s label mask, and measure the change of the mean fluorescence intensity in that mask over a time window ranging *w* frames before and after *m*. We did this for each channel, i.e., Factin, rGBD, and myosin. This provided a temporal profile over the time window [*m − w, m* + *w*] of the mean intensity associated with the event, aligned around the middle frame *m*. We repeated this procedure for all frames *m* during the event, and then for all the events. Sample profiles are shown in Fig. 3C). Thanks to the temporal alignment of the profiles, we could pool the data and calculate the averaged dynamics of the 3 markers, as shown in Fig. 3D.

## ACKNOWLEDGEMENTS

This work was supported by Swiss National Science Foundation grants 31003A163061, 51PHPO-163583, Div3 310030_185376, IZKSZ3_62195, and by a Swiss Cancer League grant KLS-4867-08-2019 to O.P., by a Human Frontier Science Program grant RGP0043/2019 to O.P. and A.C., and by a Chan Zuckerberg Initiative napari Plugin Foundation Grant 2022-252527 to M.D. and B.G.

## AUTHOR CONTRIBUTIONS

BG, OP and MD conceived the study. BG, PAG and LH performed experiments. BG and MD wrote the manuscript. BG and LB developed the Python package and the napari plugin. BG, LB, LH, and PAG analysed and visualised the data.

## Supplementary Figures

**Fig. S1:**
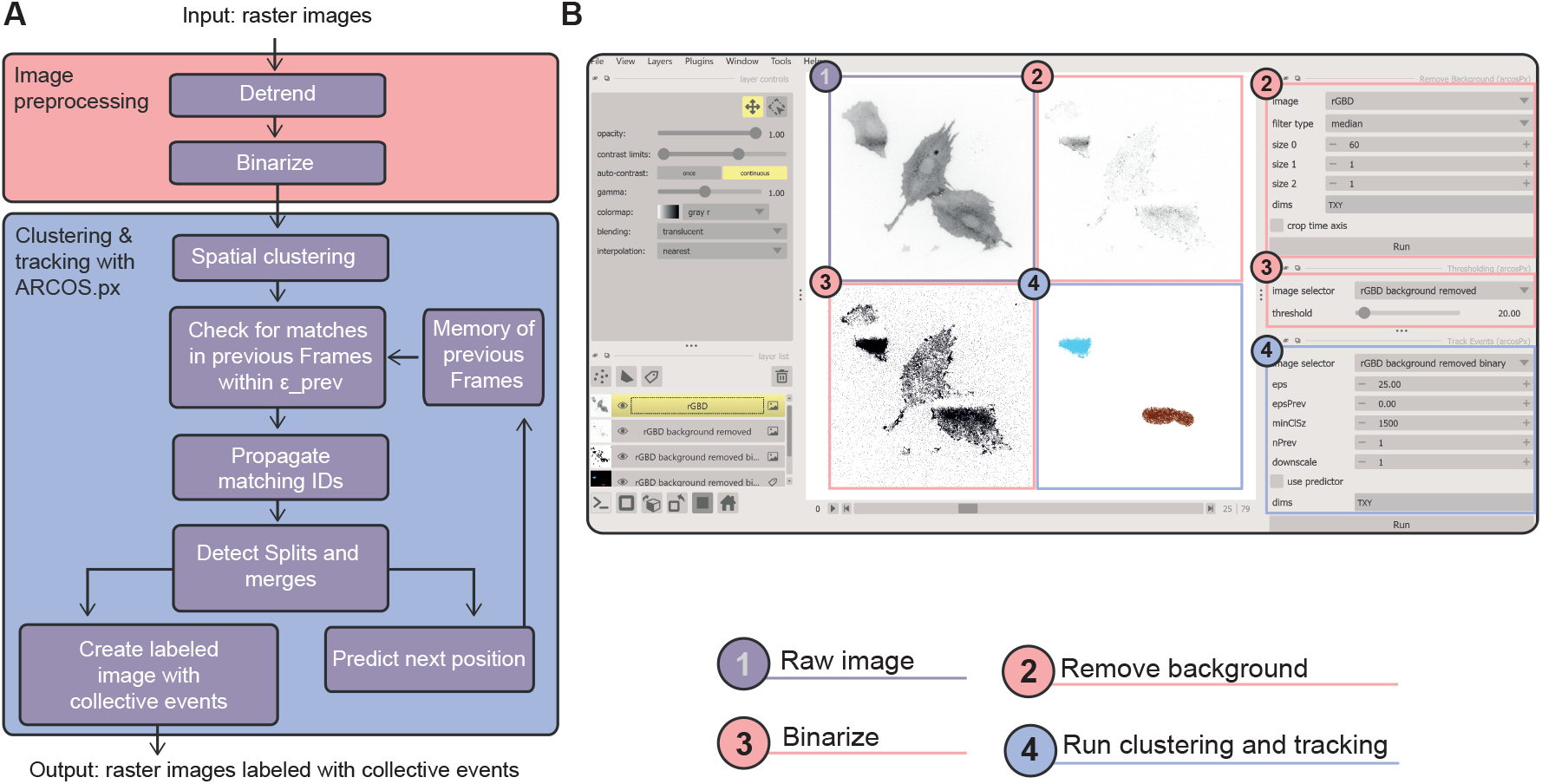
Algorithm and napari plugin. (A) Flow diagram of the ARCOS.px algorithm. (B) Screenshot of the ARCOS.px napari plugin. Circles outline the typical workflow from raw image to tracked output.

**Fig. S2:**
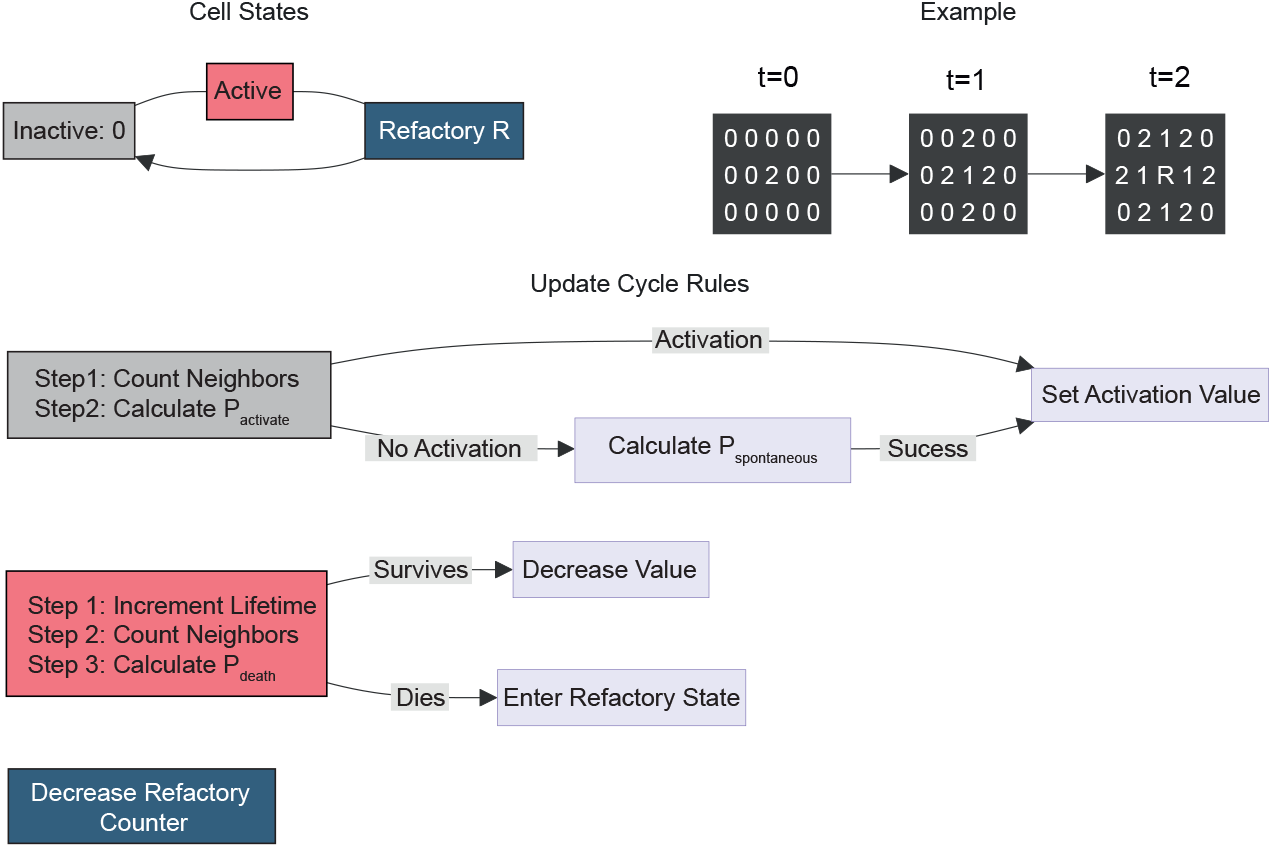
Cellular Automaton Rules. Graphical summary of the cellular automaton rules used to generate wave simulations in Fig. S3, and Video S1, Video S2, Video S3, Video S4, Video S.5. The computer code available on GitHub (23)

**Fig. S3:**
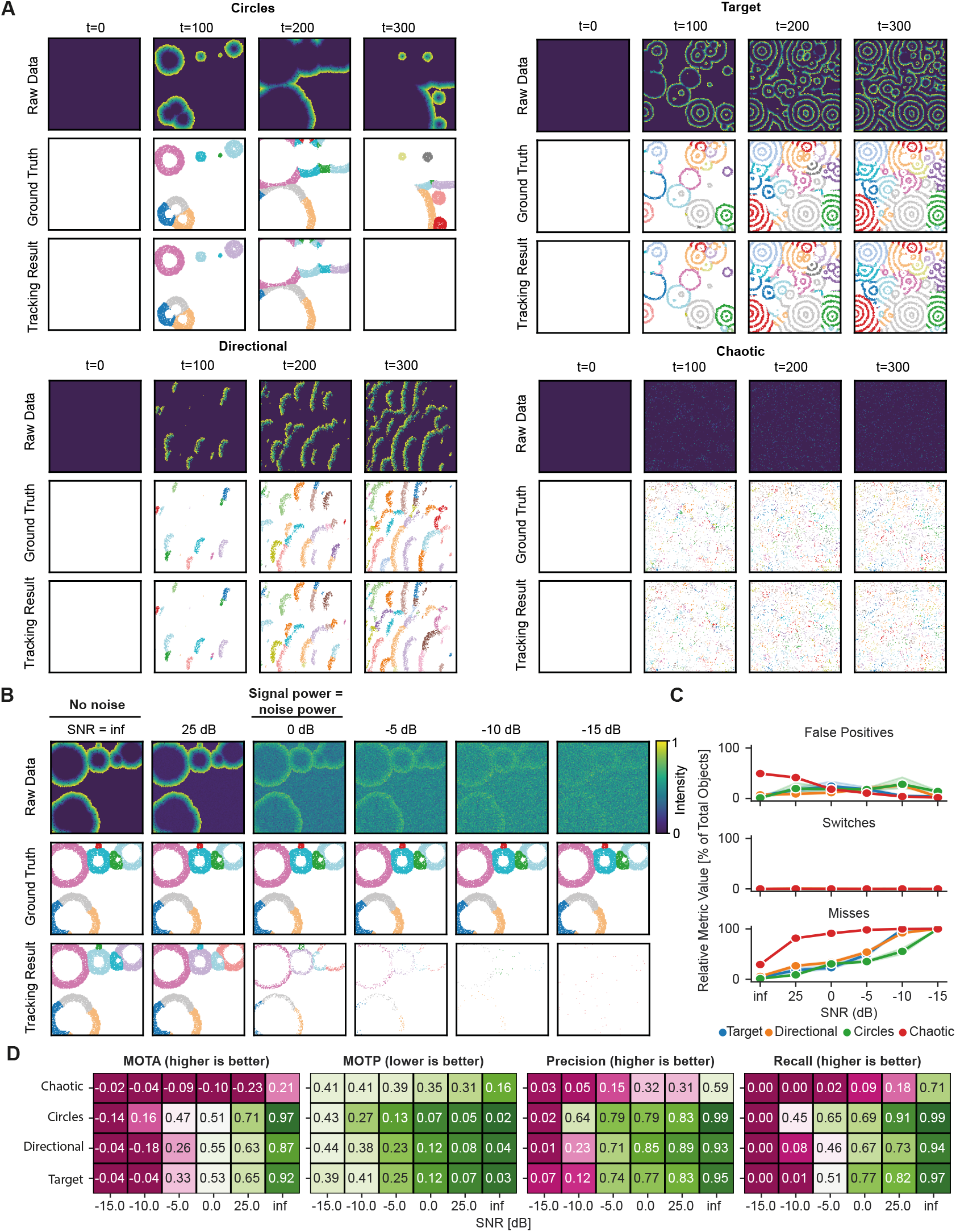
Testing the algorithm on simulated data. (A) Representative snapshots from the cellular automaton simulations of circular and directional waves, and target and chaotic patterns. Ground truth represents ID assignment by the cellular automaton. Tracking results represent raw output from ARCOS.px. (B) Circular waves with additive Gaussian noise to achieve a target signal-to-noise ratio (SNR). SNR=inf corresponds to detection without added noise. SNR=0 dB is when the signal power equals noise power. (C) Selected metrics over increasing added noise obtained from the py-motmetric python package. False positives happen when detection output has no match in the ground truth. Switches denote the number of times the existing track ID switches to a new one. Misses occur when a ground truth object is not detected in the output with an overlap threshold, *t*_*d*_ = 0.5. (D) Heatmaps for Multiple Object Tracking Accuracy (MOTA), Multiple Object Tracking Precision, Precision, and Recall of our algorithm for a given combination of pattern type and signal-to-noise ratios. For MOTA, Precision, and Recall, higher is better; for MOTP, lower is better.

**Fig. S4:**
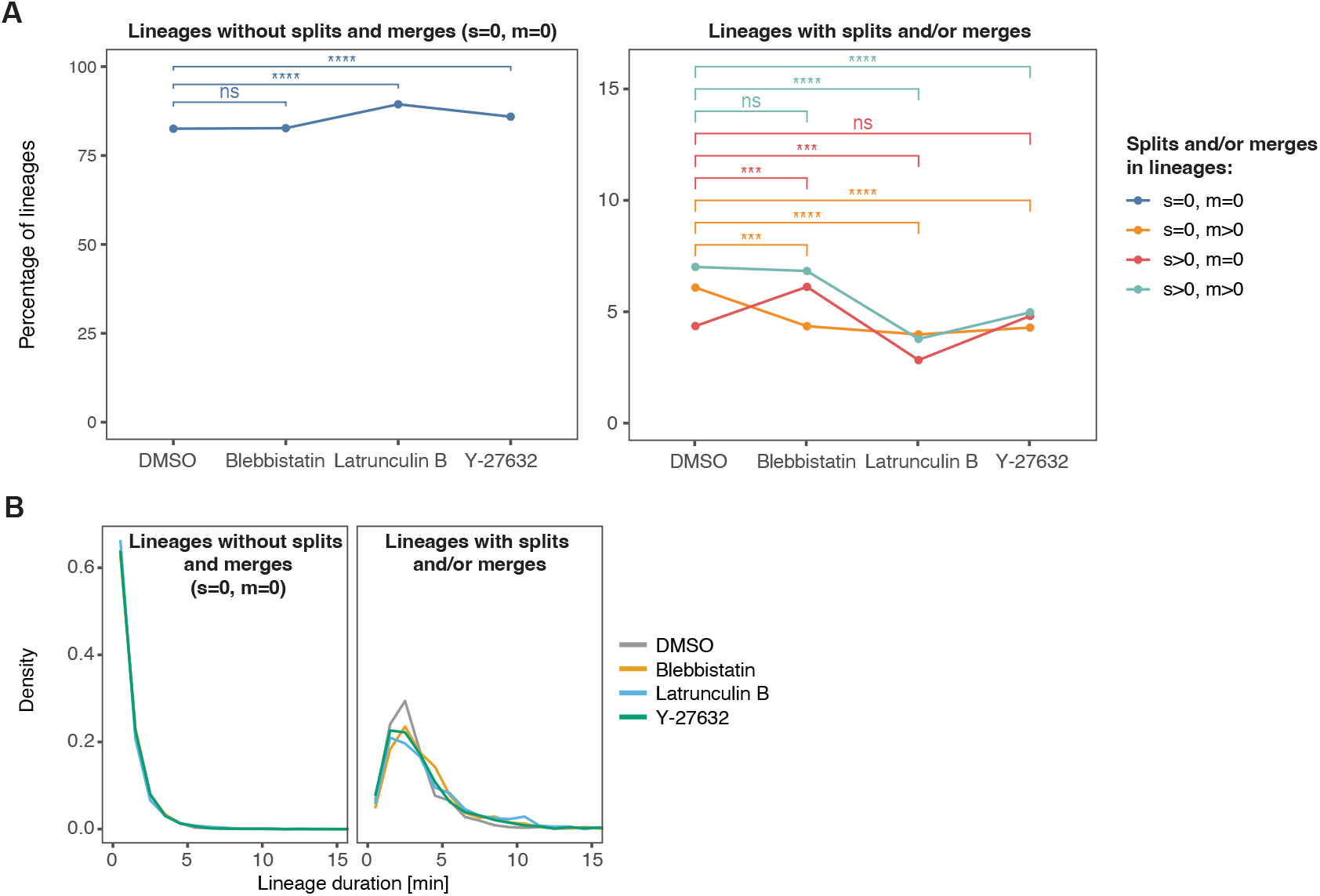
Lineage statistics in response to drug treatments. (A) The change in percentage of PLS lineages without splits and merges (s=0, m=0), without splits but with merges (s=0, m>0), with splits but without merges (s>0, m=0), with splits and merges (s>0, m>0). A two-sample test of proportions estimates the difference between groups; ****: *p ≤* 0.0001, ***: *p ≤* 0.001, **: *p ≤* 0.01, *: *p ≤* 0.05, ns: *p >* 0.05. (B) Distribution of PLS lineage duration for lineages without any splits and merges (left), and all other lineages (right).

### Supplementary Videos

**Video S1**. Detection and tracking of emergent dynamics with ARCOS.px in four types of numerically simulated pattern types: circular and directional waves, and target and chaotic patterns.

**Video S2**. Detection and tracking of emergent dynamics with ARCOS.px in numerically simulated **circular waves**. Image degradation was imitated by increasing the amount of Gaussian noise to achieve a target signal-to-noise ratio (SNR).

**Video S3**. Detection and tracking of emergent dynamics with ARCOS.px in numerically simulated **directional waves**. Image degradation was imitated by increasing the amount of Gaussian noise to achieve a target signal-to-noise ratio (SNR).

**Video S4**. Detection and tracking of emergent dynamics with ARCOS.px in numerically simulated **target patterns**. Image degradation was imitated by increasing the amount of Gaussian noise to achieve a target signal-to-noise ratio (SNR).

**Video S5**. Detection and tracking of emergent dynamics with ARCOS.px in numerically simulated **chaotic patterns**. Image degradation was imitated by increasing the amount of Gaussian noise to achieve a target signal-to-noise ratio (SNR).

**Video S6**. Optogenetic induction of a synthetic RhoA activity wave in REF52 cells expressing the RhoA activity sensor 2xrGBD-dTomato and the optogenetic actuator optoLARG. 2xrGBD-dTomato fluorescence with the optogenetically stimulated region highlighted in blue (left) is detrended (2^nd^ left), followed by its binarized representation (2^nd^ right). Binarized output is tracked with ARCOS.px and visualized in napari (right).

**Video S7**. Podosome formation in REF52 cells expressing Lifeact-mNeonGreen. Semantic segmentation is performed using convpaint (14) on actin-rich structures. The segmentation mask passed to ARCOS.px to track regions. Colours represent lineage IDs. Bounding box rectangles mark object IDs, which are assigned anew after a split or merge event.

**Video S8**. Sample crops of FOVs of podosome formation in REF52 cells in response to blebbistatin, latrunculin B, and Ycompound drug treatments. Colours represent lineage IDs.

**Video S9**. Polymerisation wave in REF52 cells. (Left panel) Raw TIRF time-lapse of REF52 fibroblasts treated with 50 ng/mL PDGF, 24h before imaging, expressing Lifeact-mNeonGreen. (Right) Lifeact-mNeonGreen TIRF time-lapse overlaid with ARCOS.px-tracked events. Colours represent lineage IDs. Bounding box rectangles mark object IDs, which are assigned anew after a split or merge event.

**Video S10**. Hole formation in the MCF10A WT epithelial monolayer in response to the doxorubicin treatment. (Left column) Mean fluorescence intensity from raw ERK-KTR (ERK activity) and FoxO-KTR (AKT activity) channels. (Middle) Holes segmented with Ilastik’s pixel classifier. Colours correspond to the probability of a pixel being part of a hole, with dark blue colours indicating 0 and bright yellow 1. (Right) Holes tracked with ARCOS.px. Colours correspond to individual holes.

### Word Counts

~~~
File: output.tex
Encoding: utf8
Sum count: 4496
Words in text: 3705
Words in headers: 43
Words outside text (captions, etc.): 689
Number of headers: 14
Number of floats/tables/figures: 6
Number of math inlines: 59
Number of math displayed: 0
Subcounts:
   text+headers+captions (#headers/#floats/#inlines/#displayed)
   34+9+0 (1/0/0/0) _top_
   13+0+0 (0/0/0/0) main
   580+1+0 (1/0/0/0) Section: Introduction
   0+1+0 (1/0/0/0) Section: Results
   388+2+147 (1/1/7/0) Subsection: Algorithm Overview
   114+1+0 (1/0/0/0) Subsection: Pre-processing
   69+1+0 (1/0/0/0) Subsection: Availability
   492+3+0 (1/0/4/0) Subsection: Testing the algorithm
   552+4+198 (1/1/25/0) Subsection: Tracking Dense Actin Structures
   385+10+138 (1/1/10/0) Subsection: Actin polymerization waves are trailed by RhoA activity and
   302+5+140 (1/1/11/0) Subsection: Tracking integrity of epithelial monolayer and
   126+1+0 (1/0/2/0) Subsection: Limitations
   650+1+0 (1/0/0/0) Section: Conclusions
   0+2+66 (1/2/0/0) Section: Supplementary Figures 0+2+0 (1/0/0/0) Section: Word Counts
~~~

---

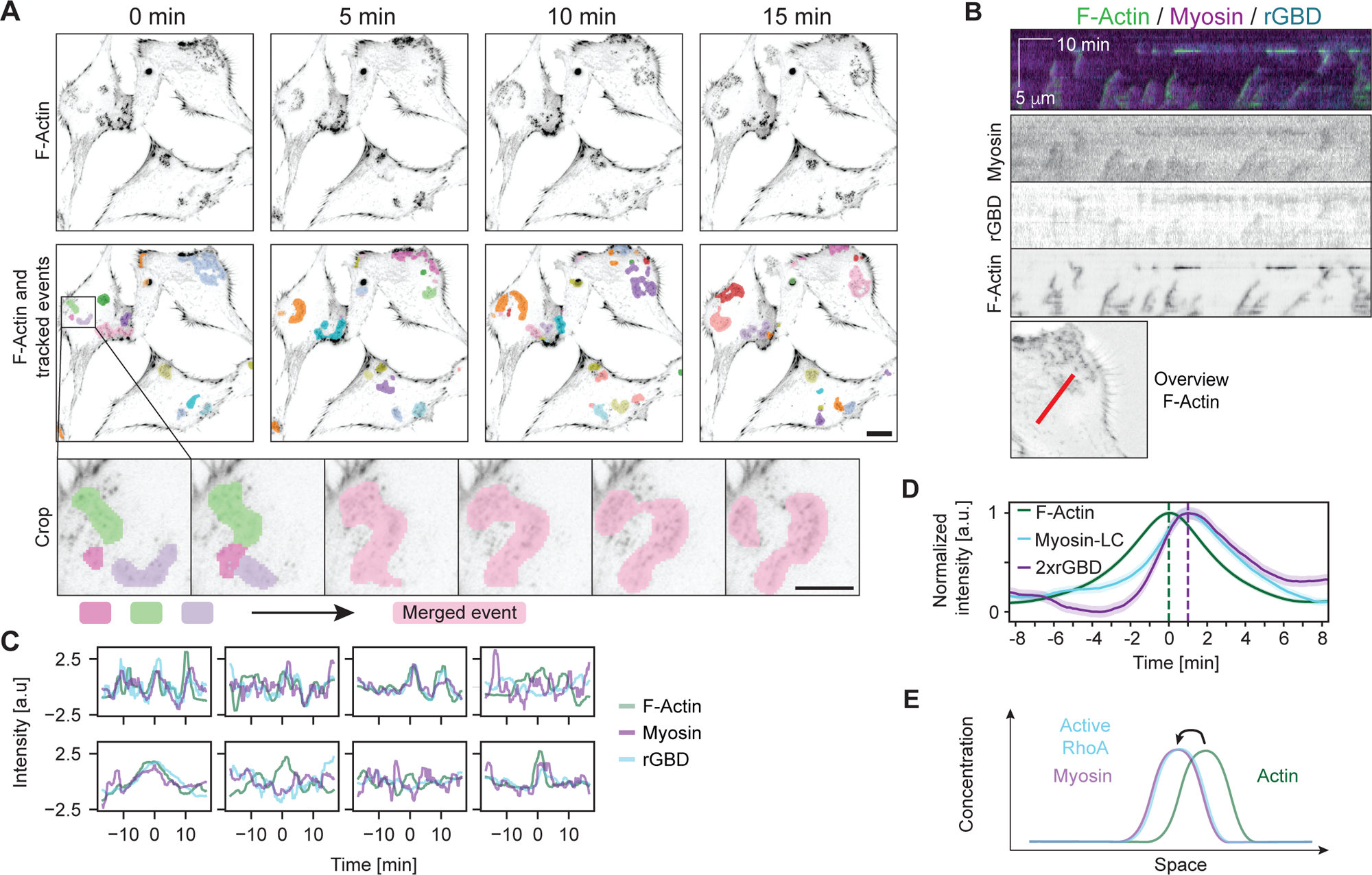

## Bibliography

1. William M. Bement, Andrew B. Goryachev, Ann L. Miller, and George von Dassow. Patterning of the cell cortex by rho gtpases. Nature Reviews Molecular Cell Biology, Jan 2024. ISSN 1471-0080. doi: 10.1038/s41580-023-00682-z.

2. Sari Tojkander, Gergana Gateva, and Pekka Lappalainen. Actin stress fibers – assembly, dynamics and biological roles. Journal of Cell Science, 125(8):1855–1864, 04 2012. ISSN 0021-9533. doi: 10.1242/jcs.098087.

3. Alex Veksler and Nir S. Gov. Calcium-actin waves and oscillations of cellular membranes. 97(6):1558–1568. ISSN 00063495. doi: 10.1016/j.bpj.2009.07.008.

4. York Posor, Wonyul Jang, and Volker Haucke. Phosphoinositides as membrane organizers. 23(12):797–816. ISSN 1471-0072, 1471-0080. doi: 10.1038/s41580-022-00490-x.

5. Yi Zhang, Geeta J. Narlikar, and Tatiana G. Kutateladze. Enzymatic reactions inside biological condensates. Journal of Molecular Biology, 433(12):166624, 2021. ISSN 0022-2836. doi: 10.1016/j.jmb.2020.08.009. Phase Separation in Biology and Disease: The Next Chapter.

6. Zsuzsanna Püspöki, Martin Storath, Daniel Sage, and Michael Unser. Transforms and Operators for Directional Bioimage Analysis: A Survey, pages 69–93. Springer International Publishing, Cham, 2016. ISBN 978-3-319-28549-8. doi: 10.1007/978-3-319-28549-8_3.

7. Erik Meijering, Oleh Dzyubachyk, and Ihor Smal. Chapter nine – methods for cell and particle tracking. In P. Michael conn, editor, Imaging and Spectroscopic Analysis of Living Cells, volume 504 of Methods in Enzymology, pages 183–200. Academic Press, 2012. doi: 10.1016/B978-0-12-391857-4.00009-4.

8. Daniel B. Allan, Thomas Caswell, Nathan C. Keim, Casper M. van der Wel, and Ruben W. Verweij. soft-matter/trackpy: v0.6.1, February 2023.

9. Benjamin Gallusser and Martin Weigert. Trackastra: Transformer-based cell tracking for live-cell microscopy. In Aleš Leonardis, Elisa Ricci, Stefan Roth, Olga Russakovsky, Torsten Sattler, and Gül Varol, editors, Computer Vision – ECCV 2024, pages 467–484, Cham, 2025. Springer Nature Switzerland. ISBN 978-3-031-73116-7. doi: 10.1007/978-3-031-73116-7_27.

10. Paolo Armando Gagliardi, Benjamin Grädel, Marc-Antoine Jacques, Lucien Hinderling, Pascal Ender, Andrew R. Cohen, Gerald Kastberger, Olivier Pertz, and Maciej Dobrzyński. Automatic detection of spatio-temporal signaling patterns in cell collectives. Journal of Cell Biology, 222(10):e202207048, 07 2023. ISSN 0021-9525. doi: 10.1083/jcb.202207048.

11. Benjamin Grädel. arcos4py: A Python Package for tracking collective signalling events. https://github.com/bgraedel/arcos4py, 2024.

12. Benjamin Grädel and Lea Brönnimann. ARCOS.px napari plugin. https://github.com/bgraedel/arcosPx-napari, 2024.

13. Stuart Berg, Dominik Kutra, Thorben Kroeger, Christoph N. Straehle, Bernhard X. Kausler, Carsten Haubold, Martin Schiegg, Janez Ales, Thorsten Beier, Markus Rudy, et al. ilastik: interactive machine learning for (bio)image analysis. Nature Methods, 16(12):1226–1232, Dec 2019. ISSN 1548-7105. doi: 10.1038/s41592-019-0582-9.

14. Lucien Hinderling, Guillaume Witz, Roman Schwob, Ana Stojiljkovic, Maciej Dobrzyński, Mykhailo Vladymyrov, Joel Frei, Benjamin Grädel, Agne Frismantiene, and Olivier Pertz. Convpaint – universal framework for interactive pixel classification using pretrained neural networks.

15. Martin Ester, Hans-Peter Kriegel, Jörg Sander, Xiaowei Xu, et al. A density-based algorithm for discovering clusters in large spatial databases with noise. In kdd, volume 96, pages 226– 231, 1996.

16. F. Pedregosa, G. Varoquaux, A. Gramfort, V. Michel, B. Thirion, O. Grisel, M. Blondel, P. Prettenhofer, R. Weiss, V. Dubourg, et al. Scikit-learn: Machine learning in Python. Journal of Machine Learning Research, 12:2825–2830, 2011.

17. Ricardo J. G. B. Campello, Davoud Moulavi, and Joerg Sander. Density-based clustering based on hierarchical density estimates. In Jian Pei, Vincent S. Tseng, Longbing Cao, Hiroshi Motoda, and Guandong Xu, editors, Advances in Knowledge Discovery and Data Mining, pages 160–172, Berlin, Heidelberg, 2013. Springer Berlin Heidelberg. ISBN 978-3-642-37456-2.

18. Thibault Séjourné, Gabriel Peyré, and François-Xavier Vialard. Unbalanced optimal transport, from theory to numerics. Version Number: 2.

19. Henri De Plaen, Pierre-François De Plaen, Johan A. K. Suykens, Marc Proesmans, Tinne Tuytelaars, and Luc Van Gool. Unbalanced optimal transport: A unified framework for object detection. Version Number: 1.

20. Rémi Flamary, Nicolas Courty, Alexandre Gramfort, Mokhtar Z. Alaya, Aurélie Boisbunon, Stanislas Chambon, Laetitia Chapel, Adrien Corenflos, Kilian Fatras, Nemo Fournier, Léo Gautheron, Nathalie T.H. Gayraud, Hicham Janati, Alain Rakotomamonjy, Ievgen Redko, Antoine Rolet, Antony Schutz, Vivien Seguy, Danica J. Sutherland, Romain Tavenard, Alexander Tong, and Titouan Vayer. Pot: Python optimal transport. Journal of Machine Learning Research, 22(78):1–8, 2021.

21. Gary Bradski. The OpenCV Library. Dr. Dobb’s Journal of Software Tools, 2000.

22. Benjamin Grädel, Lea Brönnimann, Paolo Armando Gagliardi, Lucien Hinderling, Olivier Pertz, and Maciej Dobrzyński. Data repository for Tracking Coordinated Cellular Dynamics in Time-Lapse Microscopy with ARCOS.px. https://www.ebi.ac.uk/biostudies/bioimages/studies/S-BIAD1683, 2025.

23. Maciej Dobrzyński and Benjamin Grädel. Computer code for Tracking Coordinated Cellular Dynamics in Time-Lapse Microscopy with ARCOS.px. https://github.com/dmattek/ARCOSpx-publication/, 2024.

24. Nicholas Sofroniew, Talley Lambert, Grzegorz Bokota, Juan Nunez-Iglesias, Peter Sobolewski, Andrew Sweet, Lorenzo Gaifas, Kira Evans, Alister Burt, Draga Doncila Pop, et al. napari: a multi-dimensional image viewer for Python, July 2024.

25. Anton Milan, Laura Leal-Taixe, Ian Reid, Stefan Roth, and Konrad Schindler. MOT16: A benchmark for multi-object tracking.

26. Eike K. Mahlandt, Sebastián Palacios Martínez, Janine J. G. Arts, Simon Tol, Jaap D. van Buul, and Joachim Goedhart. Opto-rhogefs: an optimized optogenetic toolbox to reversibly control rho gtpase activity on a global to subcellular scale, enabling precise control over vascular endothelial barrier strength. eLife, 12, October 2022. doi: 10.1101/2022.10.17.512253.

27. Max Heydasch, Lucien Hinderling, Jakobus van Unen, Maciej Dobrzyński, and Olivier Pertz. Gtpase activating protein dlc1 spatio-temporally regulates rho signaling. eLife, 12, June 2023. doi: 10.1101/2023.06.19.545304.

28. Eike K. Mahlandt, Janine J. G. Arts, Werner J. van der Meer, Franka H. van der Linden, Simon Tol, Jaap D. van Buul, Theodorus W. J. Gadella, and Joachim Goedhart. Visualizing endogenous rho activity with an improved localization-based, genetically encoded biosensor. Journal of Cell Science, 134(17), September 2021. ISSN 1477-9137. doi: 10.1242/jcs.258823.

29. Katrin Martin, Marco Vilela, Noo Li Jeon, Gaudenz Danuser, and Olivier Pertz. A growth factor-induced, spatially organizing cytoskeletal module enables rapid and persistent fibroblast migration. Developmental Cell, 30(6):701–716, Sep 2014. ISSN 1534-5807. doi: 10.1016/j.devcel.2014.07.022.

30. Stefan Linder, Pasquale Cervero, Robert Eddy, and John Condeelis. Mechanisms and roles of podosomes and invadopodia. Nature Reviews Molecular Cell Biology, 24(2):86–106, September 2022. ISSN 1471-0080. doi: 10.1038/s41580-022-00530-6.

31. Anna Labernadie, Anaïs Bouissou, Patrick Delobelle, Stéphanie Balor, Raphael Voituriez, Amsha Proag, Isabelle Fourquaux, Christophe Thibault, Christophe Vieu, Renaud Poincloux, Guillaume M. Charrière, and Isabelle Maridonneau-Parini. Protrusion force microscopy reveals oscillatory force generation and mechanosensing activity of human macrophage podosomes. Nature Communications, 5(1), November 2014. ISSN 2041-1723. doi: 10.1038/ncomms6343.

32. Chen Luxenburg, Sabina Winograd-Katz, Lia Addadi, and Benjamin Geiger. Involvement of actin polymerization in podosome dynamics. Journal of Cell Science, 125(7):1666–1672, 04 2012. ISSN 0021-9533. doi: 10.1242/jcs.075903.

33. Wolfgang Alt, Oana Brosteanu, Boris Hinz, and Hans Wilhelm Kaiser. Patterns of sponta-neous motility in videomicrographs of human epidermal keratinocytes (hek). Biochemistry and Cell Biology, 73(7-8):441–459, 1995. doi: 10.1139/o95-051. PMID: 8703416.

34. Michael G. Vicker. Reaction–diffusion waves of actin filament polymerization/depolymerization in dictyostelium pseudopodium extension and cell locomotion. Biophysical Chemistry, 84(2):87–98, 2000. ISSN 0301-4622. doi: 10.1016/S0301-4622(99)00146-5.

35. Kevin C. Flynn, Chi W. Pak, Alisa E. Shaw, Frank Bradke, and James R. Bamburg. Growth cone-like waves transport actin and promote axonogenesis and neurite branching. Developmental Neurobiology, 69(12):761–779, 2009. doi: 10.1002/dneu.20734.

36. Michael G. Vicker. Eukaryotic cell locomotion depends on the propagation of self-organized reaction–diffusion waves and oscillations of actin filament assembly. Experimental Cell Research, 275(1):54–66, 2002. ISSN 0014-4827. doi: 10.1006/excr.2001.5466.

37. Jun Allard and Alex Mogilner. Traveling waves in actin dynamics and cell motility. Current Opinion in Cell Biology, 25(1):107–115, 2013. ISSN 0955-0674. doi: 10.1016/j.ceb.2012.08.012. Cell architecture.

38. Naoyuki Inagaki and Hiroko Katsuno. Actin waves: Origin of cell polarization and migration? Trends in Cell Biology, 27(7):515–526, July 2017. doi: 10.1016/j.tcb.2017.02.003.

39. Julia Riedl, Alvaro H Crevenna, Kai Kessenbrock, Jerry Haochen Yu, Dorothee Neukirchen, Michal Bista, Frank Bradke, Dieter Jenne, Tad A Holak, Zena Werb, Michael Sixt, and Roland Wedlich-Soldner. Lifeact: a versatile marker to visualize F-actin. Nature Methods, 5 (7):605–607, June 2008. ISSN 1548-7105. doi: 10.1038/nmeth.1220.

40. Mutsuki Amano, Masaaki Ito, Kazushi Kimura, Yuko Fukata, Kazuyasu Chihara, Takeshi Nakano, Yoshiharu Matsuura, and Kozo Kaibuchi. Phosphorylation and activation of myosin by rho-associated kinase (rho-kinase)*. Journal of Biological Chemistry, 271(34):20246– 20249, 1996. ISSN 0021-9258. doi: 10.1074/jbc.271.34.20246.

41. Vasundhara Rao, Benjamin Grädel, Lucien Hinderling, Jakobus Van Unen, and Olivier Pertz. Feedback regulation by the rhoa-specific gef arhgef17 regulates actomyosin network disassembly. bioRxiv, 2024. doi: 10.1101/2024.08.28.610052.

42. Paolo Armando Gagliardi, Maciej Dobrzyński, Marc-Antoine Jacques, Coralie Dessauges, Pascal Ender, Yannick Blum, Robert M. Hughes, Andrew R. Cohen, and Olivier Pertz. Collective erk/akt activity waves orchestrate epithelial homeostasis by driving apoptosis-induced survival. Developmental Cell, 56(12):1712–1726.e6, Jun 2021. ISSN 1534-5807. doi: 10.1016/j.devcel.2021.05.007.

43. Ze Gong, Koen Van Den Dries, Rodrigo A. Migueles-Ramírez, Paul W. Wiseman, Alessandra Cambi, and Vivek B. Shenoy. Chemo-mechanical diffusion waves explain collective dynamics of immune cell podosomes. 14(1):2902, 05 2023. ISSN 2041-1723. doi: 10.1038/s41467-023-38598-z.

44. Charles R. Harris, K. Jarrod Millman, Stéfan J. van der Walt, Ralf Gommers, Pauli Virta-nen, David Cournapeau, Eric Wieser, Julian Taylor, Sebastian Berg, Nathaniel J. Smith, Robert Kern, Matti Picus, Stephan Hoyer, Marten H. van Kerkwijk, Matthew Brett, Allan Haldane, Jaime Fernández del Río, Mark Wiebe, Pearu Peterson, Pierre Gérard-Marchant, Kevin Sheppard, Tyler Reddy, Warren Weckesser, Hameer Abbasi, Christoph Gohlke, and Travis E. Oliphant. Array programming with NumPy. Nature, 585(7825):357–362, September 2020. doi: 10.1038/s41586-020-2649-2.

45. Pauli Virtanen, Ralf Gommers, Travis E. Oliphant, Matt Haberland, Tyler Reddy, David Cournapeau, Evgeni Burovski, Pearu Peterson, Warren Weckesser, Jonathan Bright, et al. SciPy 1.0: Fundamental Algorithms for Scientific Computing in Python. Nature Methods, 17:261– 272, 2020. doi: 10.1038/s41592-019-0686-2.

46. Stéfan van der Walt, Johannes L. Schönberger, Juan Nunez-Iglesias, François Boulogne, Joshua D. Warner, Neil Yager, Emmanuelle Gouillart, Tony Yu, and the scikit-image contributors. scikit-image: image processing in Python. PeerJ, 2:e453, 6 2014. ISSN 2167-8359. doi: 10.7717/peerj.453.

47. J. D. Hunter. Matplotlib: A 2d graphics environment. Computing in Science & Engineering, 9(3):90–95, 2007. doi: 10.1109/MCSE.2007.55.

