## Supplementary figures and images for "Tracking Coordinated Cellular Dynamics in Time-Lapse Microscopy with ARCOS.px"

### Supplemental Figure 1

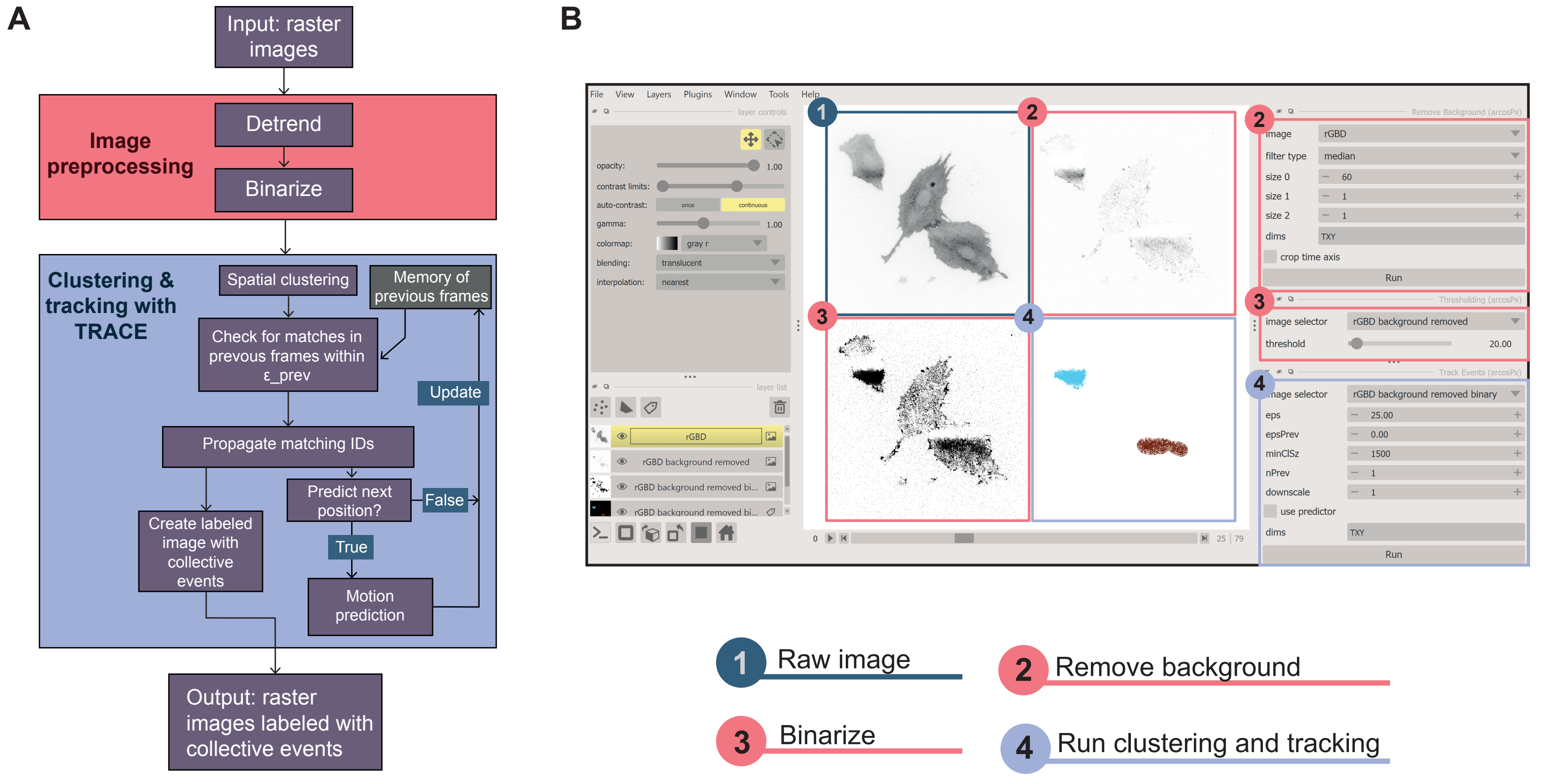

### Supplemental Figure 2

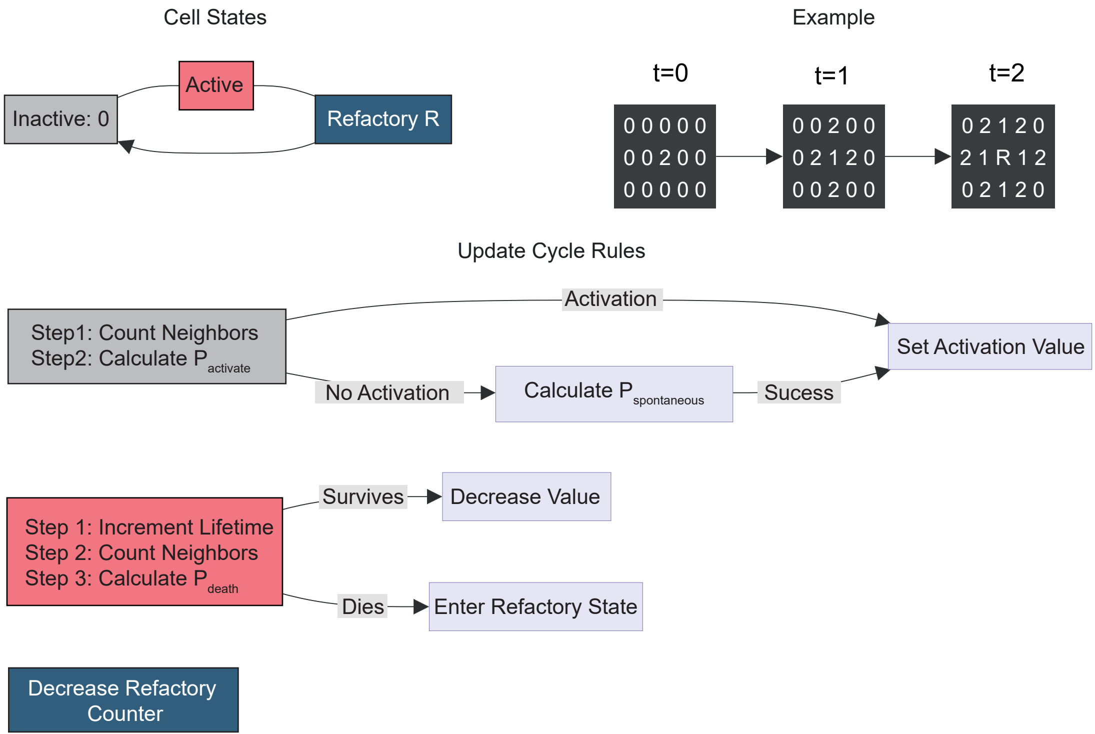

### Supplemental Figure 3

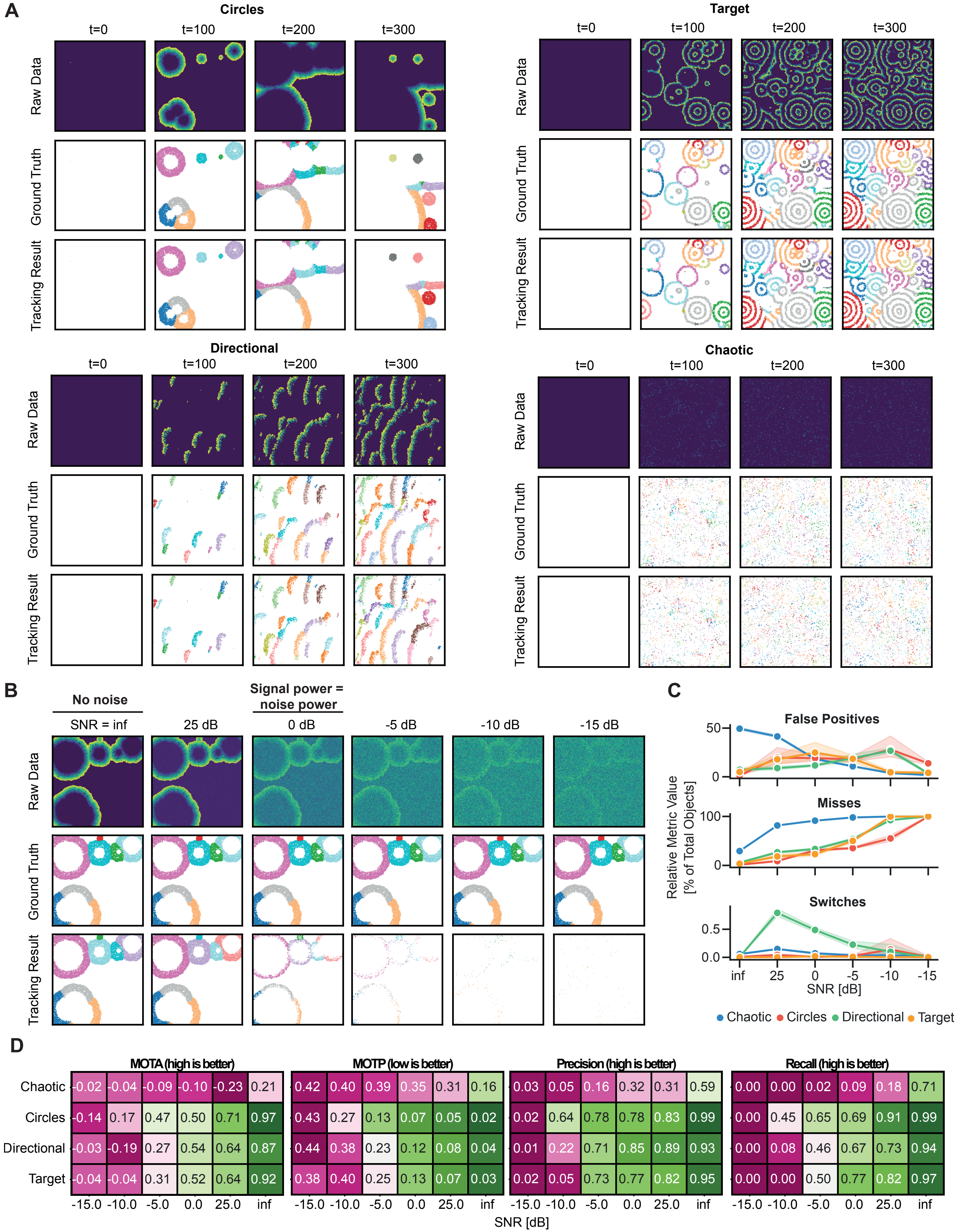

### Supplemental Figure 4

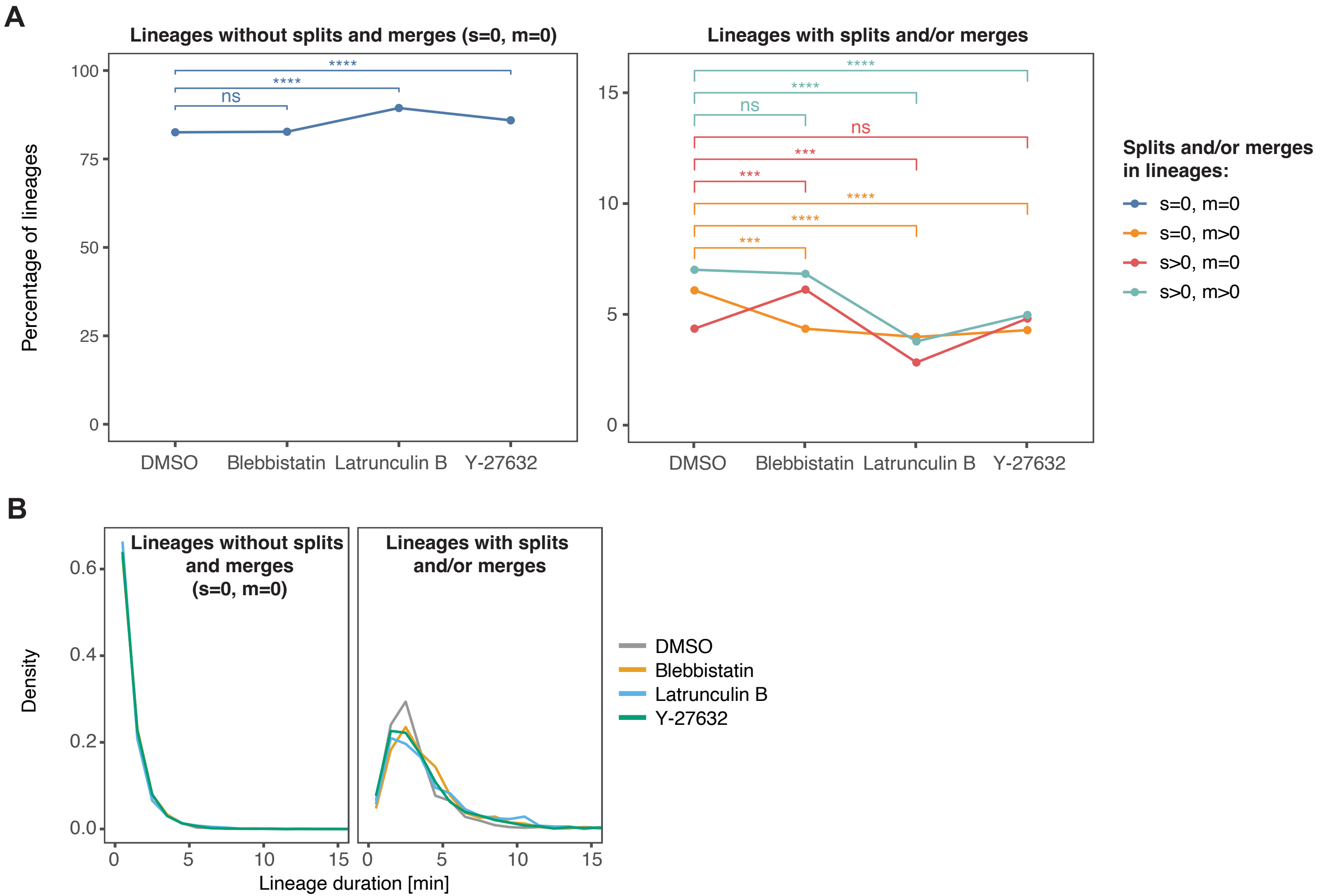
